# The structural basis for regulation of the glutathione transporter Ycf1 by regulatory domain phosphorylation

**DOI:** 10.1101/2021.06.18.449046

**Authors:** Nitesh Kumar Khandelwal, Cinthia R. Millan, Samantha I. Zangari, Samantha Avila, Dewight Williams, Tarjani M. Thaker, Thomas M. Tomasiak

**Affiliations:** Department of Chemistry and Biochemistry, University of Arizona; Tucson, AZ 85721; Department of Biochemistry and Biophysics, University of California – San Francisco, San Francisco CA, 94158; Eyring Materials Center, Arizona State University; Tempe, AZ 85287

## Abstract

Yeast Cadmium Factor-1 (Ycf1) sequesters heavy metals and glutathione into the vacuole to counter cell stress. Ycf1 belongs to the ATP binding cassette C-subfamily (ABCC) of transporters, many of which are regulated by phosphorylation on intrinsically disordered domains. The regulatory mechanism of phosphorylation is still poorly understood. Here, we report two cryo-EM structures of Ycf1 at 3.4Å and 4.0Å in distinct inward-facing open conformations capturing a previously unobserved ordered state of the intrinsically disordered regulatory domain (R-domain). R-domain phosphorylation is clearly evident and induces a topology promoting electrostatic and hydrophobic interactions with Nucleotide Binding Domain 1 (NBD1) and the lasso domain. These interactions stay constant between the structures and are related by rigid body movements of the NBD1/R-domain complex. Biochemical data further show R-domain phosphorylation reorganizes the Ycf1 architecture and is required for maximal ATPase activity. Together, we provide long-sought after insights into how R-domains control ABCC transporter activity.

## Introduction

ATP-binding cassette (ABC) transporters regulate the movement of diverse molecules to support fundamental roles of the membrane including lipid homeostasis, ion transport, and detoxification. The Yeast Cadmium Factor 1 (Ycf1) performs a similar protective function by transporting the tripeptide glutathione, which is the main non-protein thiol in cells, upon oxidation to maintain redox balance or after conjugation to toxic heavy metals such as cadmium, mercury, or lead into vacuoles^1–5^. These metals make up three of the top ten environmental toxins as defined by the CDC (#s 1,3, and 7 - https://www.atsdr.cdc.gov/spl/index.html#2019spl), marking Ycf1 as an attractive bioremediation target. Ycf1 also acts as a major redox sink and regulates redox levels in cytoplasm by sequestering glutathione after it is oxidized. In this way, Ycf1 serves a powerful protective function against metals and ROS, both of which are electrophiles that attack DNA, proteins, and the cell membrane.

Because of their essential roles in responding to cell stress, such proteins are often tightly regulated. Protein phosphorylation represents one such mechanism by which protein and transporter function can be tuned for specific cellular contexts. In general, approximately half of all human ABC transporters are phosphorylated^6^ including the medically important P-glycoprotein^7^, the Sulfonyl Urea Receptor (SUR1^8^), and the Cystic Fibrosis Transmembrane Conductance Regulator (CFTR^9^). Rapid regulation is especially important to the C-subfamily of ABC transporters (ABCC family), which includes Ycf1 and CFTR, in particular because of their roles in regulating osmotic balance, neutralizing reactive oxygen species, or combating the effects of antifungal or anticancer agents in many organisms. Often, transporter phosphoregulation is mediated through phosphorylation on regions outside of the highly structurally conserved transport machinery. Instead, phosphorylation sites arise in long, disordered loops between domains or at the termini. In ABCC transporters, such regulatory elements are evolutionary additions to the canonical ABC exporter fold composed of two transmembrane domains (TMDs) and two nucleotide binding domains (NBDs) (**Fig. 1A**). These additions include an accessory TMD called TMD0, a Lasso domain that connects TMD0 and TMD1, and a ~60-140 amino acid disordered domain termed the regulatory domain (R-domain) that connects NBD1 and TMD2^10–13^. The R-domain in particular acts as a signaling and interaction hub in ABCC transporters and contains multiple PKA, PKC, and CKII recognitions sites with otherwise little sequence conservation.

**Fig. 1.**
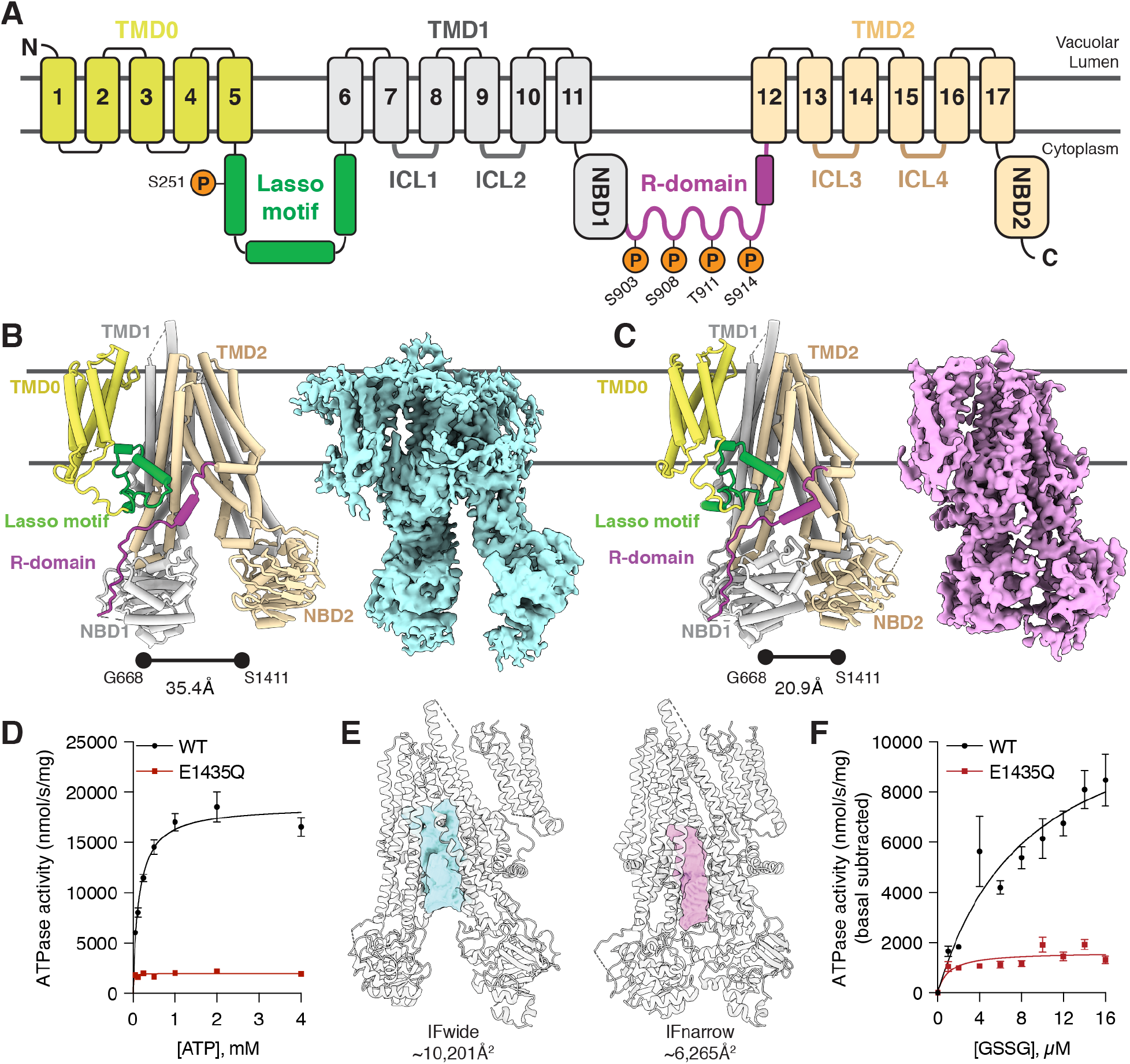
The regulatory architecture of Ycf1 determined by cryo-EM. **A.** Schematic of Ycf1 structural topology highlighting the arrangement of transmembrane helices belonging to transmembrane domain 0 (TMD0, yellow), transmembrane domain 1 (TMD1, grey), and transmembrane domain 2 (TMD2, wheat). Conserved cytoplasmic elements shown include the lasso domain (green), the R-domain (purple), nucleotide binding domains 1 and 2 (NBD1 (grey); NBD2 (wheat)) and intracellular loops 1-4 (ICL1, ICL2 (dark grey); ICL3, ICL4 (brown)). Orange spheres corresponding to conserved putative phosphorylation sites in Ycf1 are labeled with their residue numbering on the corresponding domains. Cryo-EM model (left) and map (right) of the (**B**) IFwide (cyan) and (**C**) IFnarrow (pink) conformations of E1435Q Ycf1 colored using the same scheme as shown in (**A**). **D.** ATPase activity in the presence of increasing concentrations of ATP for wild-type (WT) and the catalytically dead (E1435Q) variant of Ycf1. **E**. Volume representation of the internal cavities of IFwide (left, cyan) and IFnarrow (right, pink). **F.** ATPase activity in WT and E1435Q Ycf1 in the presence of increasing concentrations of oxidized glutathione (GSSG) and 1 mM ATP. Data are reported as stimulated rates in which the basal ATPase activity (without GSSG) was subtracted. Data shown correspond to a half-maximal effective concentration (EC_50_) of 8.67±2.6 μM for GSSG-induced stimulation of ATPase activity in WT Ycf1. Results in (**D**) and (**F**) are the mean ± the standard deviation (S.D.) for n=3 (technical triplicates).

Disruption of this regulatory function can have profound cellular effects. Mutations of R-domain phosphorylation sites lead to loss of cadmium tolerance in Ycf1^11^ and to lower open probability (but not overall open duration) and sensitivity to ATP concentration in the Ycf1 homolog CFTR^14,15^. Furthermore, mutations in 19 of the total ~130 residues of the CFTR R-domain are associated with mutants in the CFTR-linked lung disease, cystic fibrosis (www.cftr2.org). Phosphorylation can induce negative regulation as well, particularly on the lasso domain in both Ycf1^16^ and the Ycf1 functional homolog Multidrug Resistance Protein 1 (MRP1^17^). Finally, the R-domain provides a surface for protein-protein interactions, including to other transporters such as SLC26A3 and the adapter proteins 14-3-3β^18^.

Despite recent advances in ABC transporter structural biology using cryo-electron microscopy (cryo-EM), the biophysical basis of these diverse R-domain interactions is poorly understood. Barriers to these characterizations hinge on the nature of the R-domain, which is predicted to be unstructured based on low sequence conservation^19^ and experimentally confirmed with circular dichroism (CD)^19,20^ and proteolysis analysis that estimate as low as 5% helical content^20^. However, more recent NMR experiments point to discrete helical fragments in the R-domain and direct interactions with NBD1^21^. Meanwhile, cryo-EM structures of ABCC1,CFTR, and Ycf1 reveal a mostly disordered R-domain^22–24^ without assigned sequence for the phosphorylation sites, limiting our understanding of R-domain interactions. Structures of human CFTR (hCFTR) and Ycf1 have revealed density for segments of the R-domain in proximity to NBD1^25^. Unfortunately, the lack of R-domain sequence assignment in these models prohibits insights into the precise interacting regions^25^ and to date no structure of an intact R-domain with the NBDs has been determined in any state.

In this study, we have used cryo-EM to determine the 3.4Å and 4.0Å resolution structures of endogenously phosphorylated *Saccharomyces cerevisiae* Ycf1 in two inward-facing conformations: wide (IFwide) and narrow (IFnarrow). We observe a resolved section of the R-domain that is most highly conserved in the ABCC family and includes the phosphorylation sites (**Fig. 1B-C; Supplementary Table S1**). Our structures reveal the extensive interaction interface between the phosphorylated R-domain, NBD1, and the lasso-domain. Interestingly, the networks we observe are nearly constant between the two states, related only by a rigid movement of the entire R-domain/NBD1 complex relative to NBD2, with very minor changes in the positioning of the R-domain along NBD1. Biochemical investigation shows that phosphorylation induces changes in ATPase activity at the same time as it causes structural changes. Thus, we provide the structural and biochemical details of phosphorylation-dependent regulation of Ycf1.

## Results

### Cryo-EM structures of inward-facing wide and narrow states of Ycf1

In asymmetric ABC transporters, like Ycf1, only one of two NBDs (NBD2) is capable of efficient ATP hydrolysis. To enable structural studies of Ycf1 we expressed and purified two forms the *S. cerevisiae* DSY5 strain: wild-type (WT) and the NBD2 Walker B mutant E1435Q (**Fig. S1**). Grids of both WT and E1435Q Ycf1 in apo conditions were prepared as described in the methods, but only the E1435Q Ycf1 dataset yielded homogenous particles from which we successfully identified not one but two inward-facing conformations determined to 3.4Å and 4.0Å resolution (**Fig. 1A-C**). The E1435Q mutant resulted in substantially lowered ATPase activity compared to WT Ycf1 (Vmax 18633 ± 474.7 nmol/sec/mg) (**Fig 1D**), likely allowing the trapping of these conformations. Both maps were sufficiently detailed to enable modeling of an almost complete Ycf1 protein structure. The features we observe are largely consistent with other ABCC structures ^26–28^. This includes 17 transmembrane helices of TMD1 and TMD2, the accessory transmembrane domain TMD0 unique to the ABCC family, the lasso motif, both NBDs, and surprisingly a major portion of a previously uncharacterized region of the R-domain (**Fig. 1A-C**).

The two states are defined by a clear differences in the interdomain distances between the NBDs (**Fig. 1B and C**) and the TMDs that define the substrate binding cavity (**Fig. 1E**). We designated the state in which the NBDs are further apart as IFwide (**Fig. 1B**) and the state in which the NBDs are closer as IFnarrow (**Fig. 1C**). The unoccluded substrate cavities in both structures are large enough to accommodate oxidized glutathione (GSSG) (**Fig. 1E**) previously shown to be a primary physiological substrate in cells^29^. These results are consistent with our ATPase data that show GSSG indeed stimulates ATPase activity at physiologically relevant concentrations (μM concentration range) (**Fig. 1F; Fig. S1E**).

### The Ycf1 R-domain is in a phosphorylated state with an ordered regulatory domain

The most striking findings of our structures are the visualization of every regulatory element, including the R-domain (residues 901-935) phosphorylated in three positions. The R-domain is helical (residues 920-929) in the C-terminal segment and lacks secondary structure (residues 901-919 and 930-935) in the other. The C-terminal helix was key to sequence assignment and aligns with partial models from CFTR cryo-EM structures (**Fig. S7A-E,**^25^), with helical NMR assignments in CFTR^21^, and with evolutionary coupling analysis of co-evolving residues^30^ from a multisequence alignment of 13,723 homologs (**Fig. S7E**). The R-domain encircles NBD1 in both IFwide and IFnarrow (**Fig. 1B-C**) and is stabilized by extensive contacts along a basic groove of the NBD1 perimeter, intracellular loop 4, and the lasso domain (**Fig. 2A-D; Fig. 3**). This positioning differs from the positioning and sequence assignment of short, discontinuous R-domain fragments in a previous Ycf1 structure^28^.

**Fig. 2.**
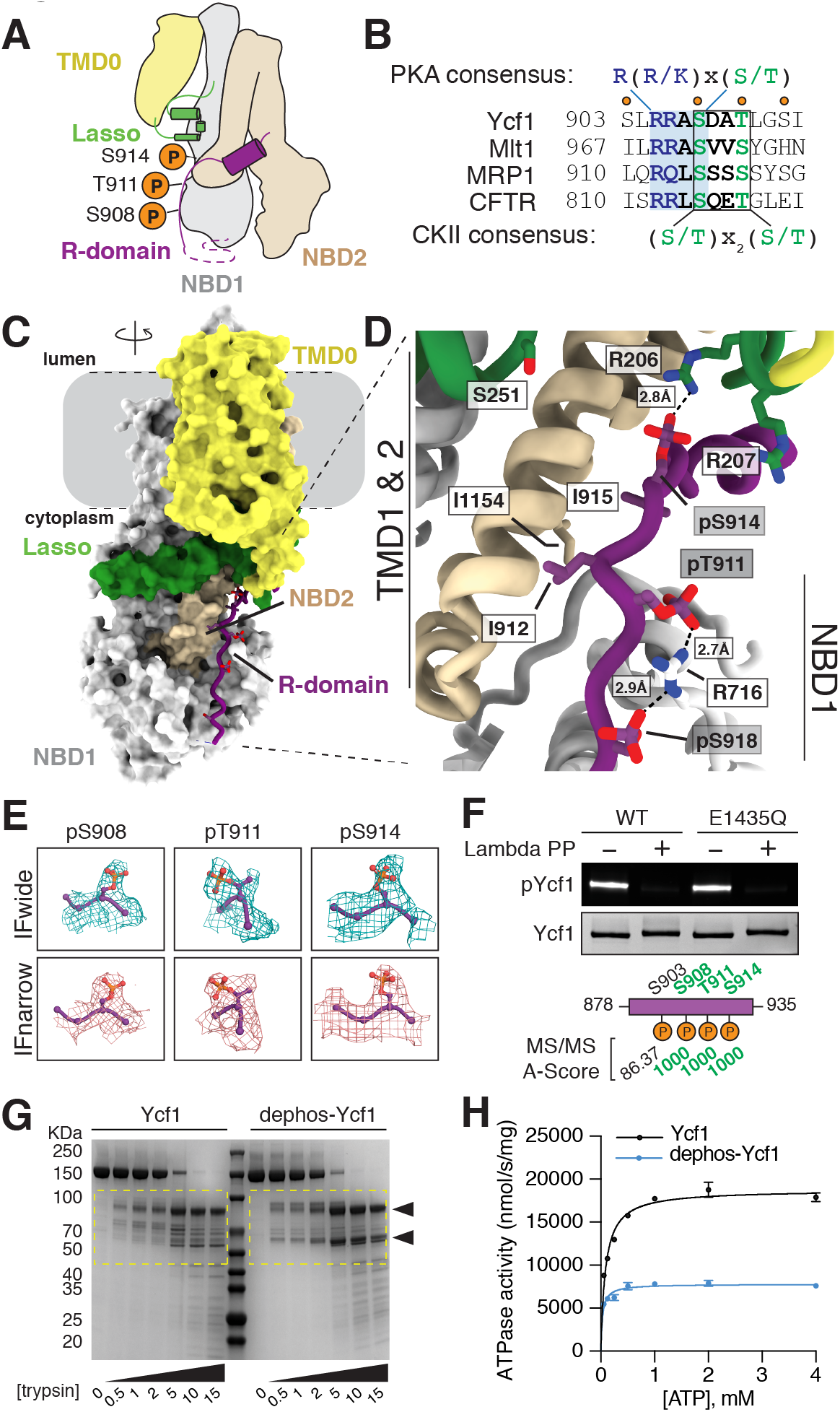
The R-domain engages Ycf1 through an extensive phosphorylation-dependent network. **A.** Putative phosphorylation sites on the Ycf1 R-domain (purple) mapped onto a cartoon representation of the Ycf1 IFnarrow cryo-EM structure. **B.** Sequence alignment of ABCC family members highlighting the consensuses sequences of putative phosphorylation motifs. Orange dots represent predicted phosphorylation sites. **C.** Surface representation of the cryo-EM model of IFnarrow Ycf1 highlighting the orientation of the R-domain along NBD1. **D**. A detailed view of the R-domain region shown in (**C**) highlighting residues contributing to the binding interface between the R-domain (purple), the lasso domain (green), and NBD1 (grey). Carbon atoms are colored consistent with domain coloring in (**C**), with oxygen (red), nitrogen (blue), and phosphate (orange) atoms colored accordingly. Dashes represent hydrogen bonds or electrostatic interactions between heavy atoms. **E.** Electron potential density of phosphorylated residues (S908, T911 and S914) observed in IFwide (cyan) and IFnarrow (pink). **F.** SDS-PAGE analysis of phosphorylation in purified samples of WT or the E1435Q variant of Ycf1 in the presence or absence of Lambda phosphatase (Lambda PP) treatment. The top gel showing phosphorylated Ycf1 (pYcf1) was stained with Pro-Q phosphoprotein gel stain (Thermo Fisher), whereas the bottom gel was visualized with Coomassie gel stain to show total Ycf1 (Ycf1). Shown below the gel are A-scores^63^ for the LC-MS/MS experiment confirming the detection of phosphorylation at the corresponding phosphorylation sites observed in the Ycf1 R-domain shown in (**D**) and highlighted in (**A-E**). **G.** SDS-PAGE analysis of proteolysis resistance in Lambda PP untreated (Ycf1) or treated (dephos-Ycf1) Ycf1 incubated with increasing concentrations of trypsin (0 – 15 μg/mL). **H.** ATPase activity of phosphorylated (Ycf1) and dephosphorylated (dephos-Ycf1). Data shown are the mean ± S.D. for n=3 (technical triplicates).

**Fig. 3.**
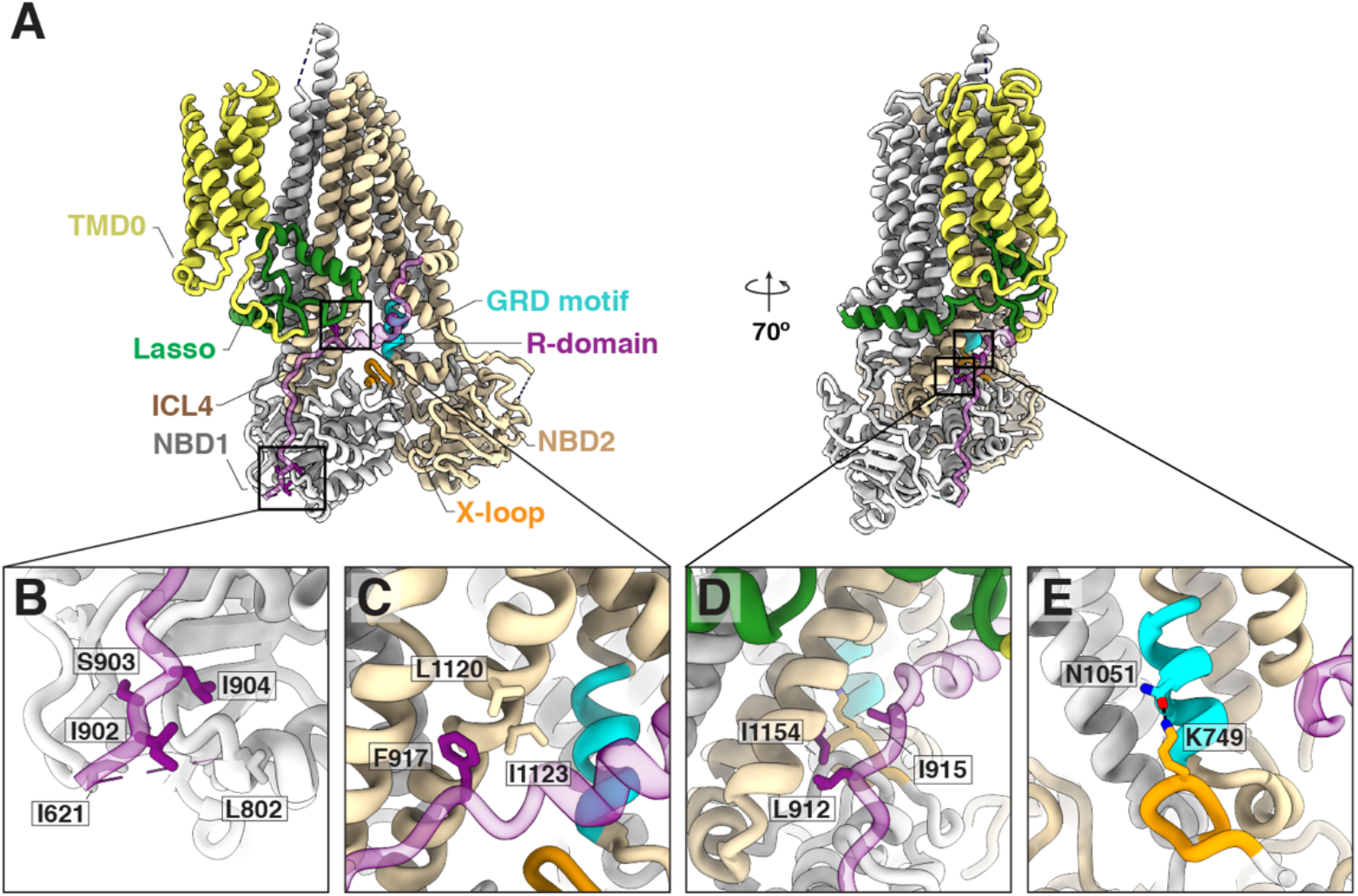
R-domain interaction network in Ycf1. **A.** Overall structure of Ycf1 in IFnarrow. **B-D**. Hydrophobic pockets along the R-domain/NBD interface. **E**. Interactions between residues of the X-loop and GRD motif in IFnarrow. The R-domain is colored purple, TMD1-NBD1 is colored grey, TMD2 and NBD2 are colored wheat, lasso domain is colored green, the X-loop is colored orange, and the GRD motif is colored cyan in all figures.

The R-domain houses several phosphorylation motifs shown to be important for regulation. We assigned phosphorylation to S908, T911, and S914 in our model based on pronounced additional electron potential density at these residues (**Fig. 2E**). Mass spectrometry and phosphostaining analysis confirm phosphorylation at these three sites in both WT and E1435Q Ycf1 (**Fig. 2F; Table S2**). These phosphates make extensive contacts with the Nε and Nη atoms in the guanidyl groups of R716 on NBD1 and R206 on the lasso domain in proximity to be either hydrogen-bonding or a strong charge-charge interactions (**Fig. 2D**). Among the phosphosites, S908 is part of a recognition motif for PKA-like kinases called the RRAS motif (or dibasic consensus sequence) and displays a similar conformation to that of a phosphomimetic peptide bound to PKA (PDB ID: 1ATP^31^). T911 also serves as part of a CKII kinase recognition motif (S/T-(x)_n_-S/T) that partially overlaps with the PKA-like kinase site (**Fig. 2B**). Overall, the resolvable range of the Ycf1 R-domain coincides with the region containing the highest sequence conservation to CFTR, which possesses a longer R-domain than Ycf1 but otherwise conserved kinase motifs (**Fig. 2B, Fig. S7E**).

### Phosphorylation of the R-domain controls dynamics and ATPase activity

The R-domain interaction with NBD1, TMDs, and lasso motif has been hypothesized to be highly dynamic and dependent on phosphorylation of the R-domain. To interrogate this effect on our reconstituted sample, we prepared dephosphorylated Ycf1 by treating it with lambda phosphatase (**Fig. 2F**) and subjecting the sample to limited proteolysis with trypsin. We observed that dephosphorylated Ycf1 substantially changes the digestion pattern in comparison to endogenously phosphorylated Ycf1(**Fig. 2G**), with a clear increase to protease accessibility in the dephosphorylated state as judged by a more complete digestion (ie loss of intermediate sized bands) in the dephosphorylated state. We reasoned that this was due to changes in the R-domain position or conformation in a phosphorylation dependent manner similar to CFTR^25^ but note that the phosphates themselves may block protease accessibility to some sites as well.

To investigate the functional impact of these structural changes due to dephosphorylation, we interrogated changes in ATPase activity. ATPase assays show a drastic activity reduction in dephosphorylated Ycf1 (**Fig. 2H**), suggesting that the interactions of phosphorylated S908, T911, and S914 are major drivers of an architecture that facilitates ATP hydrolysis. This is consistent with cell viability data from *S. cerevisiae* cells where S908 and T911 Ycf1 mutants show deficient cadmium detoxification, resulting in cell death^11^.

### Tightly embedded hydrophobic surfaces stabilize the R-domain NBD interaction

Together, the phosphorylation sites form a scaffold that engages NBD1 and the lasso motif along a single axis to connect NBD1 and the TMDs (**Fig. 3A**). The electrostatic interactions are supported by a substantial burying of hydrophobic surfaces at 3 sites (site 1: I621 - L802 - I902 - I904 (**Fig. 3B**); site 2: F917 - I1123 - L1120 (**Fig. 3C**); and site 3: L912 - I915 - I1154 (**Fig. 3D**)). These interactions are poised to provide a large thermodynamic driving force of R-domain binding and stabilize the R-domain/NBD1 interface, burying ~1530 Å^2^ and ~1338 Å^2^ of surface area along the entire R-domain in IFnarrow and IFwide, respectively (~523Å^2^ (IFnarrow) and ~480Å^2^ (IFwide) specifically between NBD1 and the R-domain) (**Fig. 2C**). In addition, this region overlaps with known allosteric motifs, including the x-loop and the GRD motif, both of which also contact each other in IFnarrow (**Fig. 3E**).

### Location of an inhibitory phosphosite at S251 in proximity to R-domain phosphosites

The lasso domain region adjacent to the R-domain also contains a conserved phosphorylation site that negatively regulates transport, S251 (T249 in the Ycf1 homolog MRP1)^16,17^. Though we do not observe phosphorylation in our structure at this position, S251 is accessible to the surface and poised to disrupt R-domain interactions through electrostatic repulsion of phosphate groups in the R-domain (**Fig. 2D**). Thus, our structures provide a plausible explanation of the negative impact of the lasso domain phosphorylation on Ycf1 function.

### The two structures of Ycf1 show different NBD orientation with a single-occluded ATP site in IFnarrow

These observed R-domain - NBD1 - lasso contacts are consistent between IFwide and IFnarrow with minimal changes between these domains (**Fig. 4A**). Instead, a rearrangement within the R-domain correlates with overall movements in the structure, including a change in angle between residues G913, G918 and G932 of the R-domain from 130° (IFwide) to 137.5° (IFnarrow) (**Fig. 4B**). The glycines G918 and G932 cap the helical R-domain region and act as a hinge for the movement of this structured region between the transition from IFwide to IFnarrow.

**Fig. 4.**
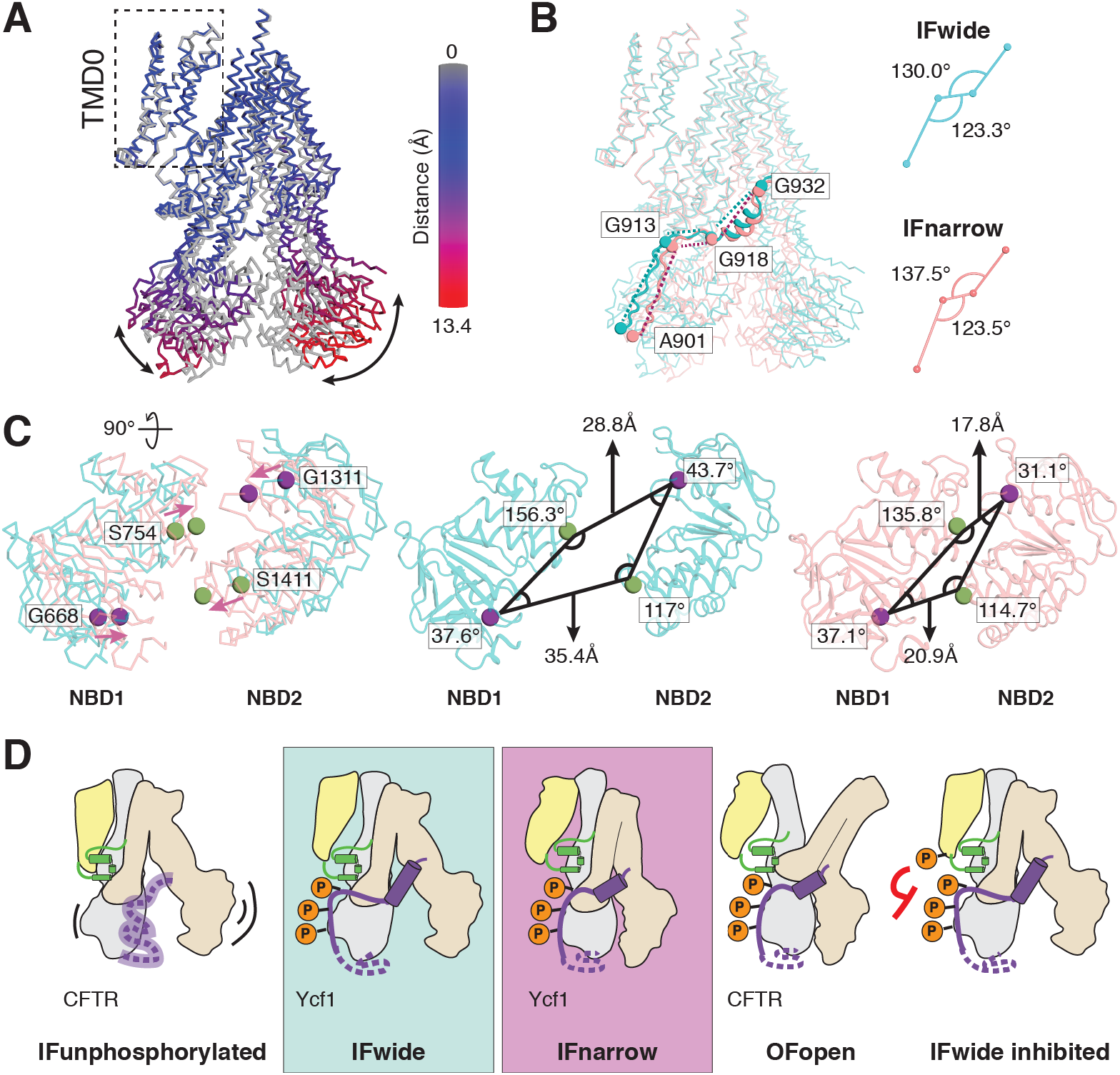
Proposed model for catalytic control in Ycf1 relies on transitions of the rigid R-domain architecture from IFwide to IFnarrow. **A.** Overlay of IFnarrow (grey) and IFwide states of Ycf1, in which IFwide is colored by RMSD calculated on a per residue basis from structural alignment between TMD0 of each state. **B.** Comparison of the R-domain geometry and differences observed for the intradomain angles between flexible residues forming hinge regions in the R-domains in each state. **C.** Superposed NBDs (left panel) from IFwide (cyan ribbon) and IFnarrow (pink ribbon) highlighting the rearrangement in the relative positions of conserved residues of the signature motif (S754 and S1411, green sphere) and the walker A motif (G668 and G1311, purple spheres) in each state. The specific interdomain angles formed by these sites in their respective states are shown in the middle (IFwide) and right (IFnarrow) panels. **D.** Proposed model for organization and regulation of the R-domain through transport in light of the IFwide and IFnarrow Ycf1 structures and in the context of previously published CFTR structures (IFunphosphorylated (PDB ID 5uak^26^); OFopen (PDB ID 6msm^25^). Domains are colored yellow for TMD0, gray for TMD1 and NBD1, wheat for TMD2 and NBD2, green for the Lasso domain, and purple for the R-domain.

In contrast to the interaction sites themselves, the major structural consequences of the IFwide to IFnarrow transition are induced at the ATP binding sites. These sites not only move closer in IFnarrow but also rotate to occlude the non-functional ATP binding site in NBD1 (**Fig. 4C**). Notably, the second site in NBD2 (active) is open for ATP binding. NBD2 is the less ordered of the two NBDs as inferred from relatively weaker cryo-EM density in this region in both states. This disorder is a predominant feature also noted for a third, less well-resolved state that we call IFtransition (**Fig. S3C**) which was insufficiently detailed to enable model building. Concurrently, several coupling elements that connect the NBDs and TMDs, including the NBD1 X-loop and GRD motif^32^, are brought into contact upon transition to IFnarrow (**Fig. 3E**). Overall, these architectural changes resemble changes in substrate-bound and substrate-free states of MRP1 (**Fig. S8A-F**)^27,33^ and point to a model where Ycf1 R-domain phosphorylation helps stabilize NBD1 to recruit other domains into a conformation (IFnarrow) suitable for ATP binding and initiation of transport. We propose this yields a primed, active state ready for transport.

## Discussion

The structures of Ycf1 determined here reveal previously unobserved insights in the ABCC family of ABC transporters that inform on how they are regulated. This includes an inward-open conformation with an intact and endogenously phosphorylated R-domain tightly bound to NBD1. This provides a more complete model than previous ABCC structures, which resolve small fragments of the R-domain, do not resolve the phosphorylation sites, and do not assign the sequence^28,34,35^. The presence of an ordered R-domain position poses a surprise, since many alternate models exist for the R-domain mechanism including a model where it is completely disordered or where it resides only between NBD1 and NBD2 in a semi-ordered state.

The findings in this work support decades of biochemical, cellular, and clinical data on the consequences of R-domain interactions. The multiple phosphate binding interactions to a common surface along NBD1 and the lasso domain explain the well-known robustness to loss of any single phosphorylation site on the R-domain^9,11^ and the requirement for an intact N-terminus containing the lasso domain of Ycf1 for activity^36^. The extensive R-domain and NBD1 interactions increase the ATPase rate in a manner consistent with CFTR, where R-domain phosphorylation potentiates ATP binding and hydrolysis but otherwise does not induce activity alone^37^. Our structures also are consistent with the visible R-domain density in CFTR and polyalanine models built into this density (**Fig. S7A-D**).

The position of the R-domain informs on the phosphorylation activation mechanism and provides clarification on the competing models that exist. In the most widely held model, CFTR activation is proposed to depend on release of the R-domain positioned between NBD1 and NBD2 upon phosphorylation, which induces a completely unbound and disordered state^26^. Our findings here do not support this model. Instead, our results suggest that the extensive interactions between the phosphorylated R-domain and the rest of Ycf1 represent a stimulatory R-domain architecture. However, our structures do not rule out an inhibitory role for the unphosphorylated state (**Fig. 4D**). Based on our functional data, we propose a model where the R-domain transitions from unbound/disordered to bound/ordered upon phosphorylation, residing either between the NBDs when dephosphorylated^38^ or near NBD1 but loosely associated. Phosphorylation along the R-domain drives association with the periphery of NBD1 (**Fig. 2C**) that is reflected as both increased ATPase activity and R-domain stability (**Fig. 2G-H**). The clustering of the R-domain and lasso domain also provides a plausible explanation for the inhibitory role of S251 phosphorylation, which is poised to disrupt the R-domain interaction network by charge repulsion (IFwide inhibited - **Fig. 4D**). In conclusion, R-domain interactions lend a regulatory effect analogous to that of a transmission “clutch” that only engages a motor but otherwise does not perform work itself.

These results also rationalize key clinically relevant findings related to other ABCC family members. The extensive electrostatic, hydrogen bonding, and hydrophobic interactions on the R-domain explain how mutations distal from phosphorylation sites may nevertheless contribute to dysfunction. Additionally, the site equivalent to the most widespread cystic fibrosis linked mutant (CFTR delF506), Ycf1 F713, is located in NBD1 directly adjacent to the phosphorylation sites of the R-domain and is coordinated by cation-pi interactions with two conserved arginine side chains, R765 and R1150 (**Fig. S7H**). Finally, the binding site of lumacaftor, a clinically used corrector that stabilizes unfolded CFTR, binds in the outer perimeter of NBD1 near the R-domain binding site ^39^. The high convergence of several important effects near one region where the R-domain, NBD1, and lasso domain come together suggest the general importance of the R-domain interactions observed in our structures.

In summary, our structures provide mechanistic insights about post-translational modification (phosphorylation) on an important class of transporters, the ABCC family. The presence of a single defined binding site for the conserved part of the R-domain confirms an anticipated but elusive binding site and may provide implications for several other types of transporters where such long phosphorylated unstructured domains or loops are widespread. Such a defined binding site could also provide a basis for allosteric modulation of transporter function in a diverse array of settings.

## Methods

### Cloning, expression, and purification

The *S. cerevisiae YCF1* (Yeast Cadmium Factor 1) gene was codon-optimized and cloned into the p423_GAL1 yeast expression vector as an N-terminal Flag (DYKDDDDK) and C-terminal decahistidine (10X His) tagged fusion protein (GenScript) (**Fig. S1A**). The E1435Q mutant was generated by site-directed mutagenesis (forward primer: 5’-CTTGGTTTTGGATCAAGCTACAGCTGCAG-3’; reverse primer 5’-CTGCAGCTGTAGCTTGATCCAAAACCAAG-3’) and verified by sequencing (Elim Biopharmaceuticals, Inc).

For protein expression, the *S. cerevisiae* strain DSY-5^40^ (Genotype MATa leu2 trp1 ura3-52 his3 pep4 prb1) was transformed with the Ycf1 expression construct and a 50 mL primary culture grown for 24 hours at 30°C with shaking at 200 rpm in SC-His media (0.67% w/v yeast nitrogen base without amino acids, 2% w/v glucose, and 0.08% w/v amino acid dropout mix without histidine). A secondary 750 mL culture of SC-His media was inoculated with 2% of the primary culture (15 mL) and grown under the same growth conditions for an additional 24 hours prior to induction by adding YPG media (1% w/v yeast extract, 1.5% w/v peptone, and 2% w/v galactose final concentration) from a 4X YPG media stock. The culture was grown for an additional 16 hours at 30°C prior to harvesting by centrifugation at 5000xg for 30 minutes at 4°C.

For protein purification, harvested cells were resuspended with ice-cold lysis buffer (50 mM Tris-Cl pH 8.0, 300 mM NaCl, and cOmplete, EDTA-free protease inhibitor cocktail tablets (Roche)) at a ratio of 3.2 mL/g of cell pellet. Resuspended cells were lysed on ice by bead beating with 0.5 mm glass beads for 8 cycles consisting of 45 seconds of beating, with 5 minutes between cycles. Lysates were collected by vacuum filtration through a coffee filter and membranes harvested by ultracentrifugation at 112,967xg for 1.5 hours prior to storage at −80°C. Membranes were solubilized in resuspension buffer (50 mM Tris-Cl pH 7.0, 300 mM NaCl, 0.5% 2,2-didecylpropane-1,3-bis-β-D-maltopyranoside (LMNG)/0.05% cholesteryl hemisuccinate (CHS) supplemented with protease inhibitor as described above) at a ratio of 15 mL/g of membrane at 4°C for 4 hours. Solubilized membranes were clarified by centrifugation at 34,155xg for 30 min at 4°C. The clarified supernatant was filtered through a 0.4 μM filter to remove the insoluble fraction and supplemented with 30 mM Imidazole pH 7.0 immediately before loading at a flow rate of 2 mL/min onto a 5 mL Ni-NTA immobilized metal affinity chromatography (IMAC) column (Bio-Rad) equilibrated in Buffer A (50 mM Tris-Cl, 300 mM NaCl, 0.01% LMNG/0.001% CHS, pH 7.0).

Following loading, the column was washed with 10 column volumes (CV) of Buffer A to remove nonspecifically bound proteins then followed by a gradient of Buffer B (50 mM Tris-Cl, 300 mM NaCl, 500 mM Imidazole 0.01% LMNG/0.001% CHS, pH 7.0) consisting of the following step sizes: 6% (10CV), 10% (2CV), 16% (2 CV), and 24% (2CV). Protein was eluted with 4CV of 60% buffer B and immediately diluted 10-fold with Buffer A prior to concentration and 3 rounds of buffer exchange to remove excess imidazole by centrifugation at 4000 rpm at 4°C in 100 kDa cutoff concentrators (Amicon). Concentrated, buffer exchanged sample was lastly purified by size exclusion chromatography (SEC) at 4°C by injecting sample onto a Superose 6 Increase 10/300 GL column (GE Healthcare) equilibrated in SEC buffer (50 mM Tris, 300 mM NaCl, pH 7.0) supplemented with either 0.01% LMNG/0.001%CHS or 0.06% digitonin and immediately used for biochemical assay or cryo-EM grid preparation following quantification by BCA Assay (Pierce).

### Cryo-EM grid preparation and data acquisition

Cryo-EM grids for wild-type (WT) and E1435Q Ycf1 were similarly prepared. Immediately following SEC purification, 5 μL of concentrated WT (15.2 mg/mL) or E1435Q (5.94 mg/mL) Ycf1 sample was applied to a CFlat-1.2/1.3-4C-T (WT Ycf1) or QF-1.2/1.3-4Au grid (E1435Q Ycf1) purchased from Electron Microscopy Sciences. Grids were placed inside of a Leica EM GP2 equilibrated to 10°C and 80% humidity. Following a 10s incubation, the side of the grid to which sample was applied was blotted on Whatman 1 paper (8s for WT; 3.5s for E1435Q), then immediately plunge frozen in liquid ethane equilibrated to −185°C. A total of 3,262 movies were captured for WT Ycf1 using Serial EM software on a Titan Krios at 300 kV equipped with a K2 Summit detector (Gatan) at Arizona State University (ASU). For E1435Q, 8,499 movies were captured on a Titan Krios at 300 kV equipped with a K3 Summit detector (Gatan) at the Pacific Northwest Center for Cryo-EM (PNCC). Movies of both WT and E1435Q Ycf1 samples were collected at 22,500X magnification with automated super resolution mode and defocus ranges of −0.5-2.8 μm (WT Ycf1) and −0.9 to −2.1 μm (E1435Q Ycf1). Movie frames for WT Ycf1 contained 40 frames with a per frame exposure of 1.4 electrons /Å^2^ (~56 electrons /Å^2^ total dose). Movies of E1435Q Ycf1 contained 60 frames with a per frame exposure of 0.9 electrons /Å^2^ dose rate (~54 electrons /Å^2^ total dose).

### Cryo-EM data processing

The Ycf1 E1435Q dataset was processed in RELION (3.0 and 3.1)^41^ and cisTEM^42^. Drift correction was performed using MotionCor2^43^ to generate an image stack with a pixel size of 1.031 Å/pixel. The contrast transfer function (CTF) was estimated for dose-weighted micrographs using CTFFIND4.1 prior to particle picking using a reconstruction of WT Ycf1 (**Fig. S2**)^44^. Manual particle picking was performed on a subset of micrographs belonging to the WT Ycf1 dataset and subject to reference free 2D Classification to generate references for automated particle picking. Following ab-initio 3D map generation and several rounds of 3D classification and 3D refinement in RELION, a ~6.0 Å resolution map of WT Ycf1 was obtained (**Fig. S2**), low-pass filtered to 20Å resolution, and used as a reference for automatic particle picking in RELION in the E1435Q mutant dataset using a 15° degree angular search (**Fig. S3**). A total 2,159,582 particles were automatically picked, extracted with 4X binning resulting in a box size of 440 pixels with 4.124 Å/pixel. Multiple rounds of 2D classification were performed to remove bad particles resulting in 1,626,297 particles subject to 3D analysis in RELION following extraction with 2X binning and a box size of 300 pixels with 2.062 Å/pixel. The two major classes corresponding to two distinct states from the second round of 3D Classification were extracted with a 300 pixel box size at the full pixel size of 1.031 Å/pixel and subjected to iterative rounds of CTF refinement, Bayesian polishing, and postprocessing in RELION. To reduce alignment bias due to the presence of the detergent micelle, SIDESPLITTER refinement^45^ was implemented in later stages of 3D Refinement in RELION. A final round of alignment-free 3D Classification to remove structural heterogeneity revealed the IFnarrow and IFwide states, as well as a third state (IFtransition) in which NBD2 was poorly resolved. Maps from RELION were further refined in cisTEM and used for manual model building. ResMap was used for local resolution estimation performed on cisTEM maps ^46^. A summary of the data processing workflow and final EM map quality is reported in **Figs. S2-5**.

### Model building and refinement

An initial model of Ycf1 was built using the SWISS-MODEL server^47^ and 6jb1 as a template^48^. Manual model building was performed in COOT^49^. Iterative cycles of real-space refinement and analysis were performed in Phenix^50^ and CCP-EM modules^51,52^ were used throughout the structure building process for map sharpening/blurring and for structure analysis. Secondary structure restraints were used extensively, and model building was guided by evolutionary couplings analysis (**Fig. 7F-G**). Modrefiner was used in the early stages of refinement to help assign secondary structure and correct geometry^53^. Isolde was used to optimize model to map fit and to improve geometry^54^. Molprobity was used extensively through Phenix and through a dedicated web service to optimize geometry^55^. To maintain proper geometry, starting model restraints and harmonic restraints were used extensively in Phenix. Analysis of the Ycf1 substrate binding cavity volume was performed using the 3V server^56^. In this calculation, the Ycf1 NBDs were excluded in order to obtain information restricted to cavity volumes in the TMDs. A probe size of 2.5Å was used as was previously used in assessment of a bacterial glutathione transporter^57^.

### Evolutionary coupling analysis of Ycf1

Evolutionary couplings analysis was performed using the Evcouplings package downloaded from https://github.com/debbiemarkslab/EVcouplings. Sequences were chosen automatically from the JackHMMR^58^ protocol supplied with the Evcouplings suite using the Uniref90 database. A total of 13,723 sequences were identified and coupling scores calculated with PLMC^30^. HH-suite was used to cluster similar sequences at an 80% cutoff that would otherwise skew the distribution of homologs^59^. Scores with a cutoff probability of ≥0.99% were used to identify positive interactions.

### ATPase activity assays

For evaluating ATPase activity, WT and E1435Q Ycf1 were expressed and purified as described above in buffer containing 0.01% LMNG and 0.001% CHS. ATPase rates were determined at 30°C using an enzyme-coupled assay previously described with slight modification^60^. Each reaction consisted of 75 μL volumes containing 6.45 μg of protein in a reaction mix of 20 mM Tris-HCl pH 7.0, 10 mM MgCl_2_, 1 mM PEP, 55.7 / 78.03 U/mL PK/LDH, 0.3 mg/mL NADH and ATP at varying concentrations. Following the addition of ATP, the initial rate of NADH consumption was monitored by measuring the absorbance every minute at 340 nM for 30-45 min on a Synergy Neo2 Multi-mode Microplate Reader (BioTek). Nonlinear regression analysis of data fit with the Michaelis-Menten equation in GraphPad Prism 9 was used to generate kinetic parameters for at least three technical replicates.

Substrate-stimulated ATPase activity was measured as described above and in the presence of varying concentrations of oxidized glutathione (GSSG) (**Fig. 1F; Fig. S1E**). The concentration of ATP in these experiments was held constant at 1 mM. Data for the basal ATPase activity in the absence of GSSG was subtracted from the substrate-stimulated data prior to fitting, as was performed in the study of ABCG2^61^. Data were fit using nonlinear regression in GraphPad Prism 9 to derive EC50 and Vmax values.

### Characterization of Ycf1 phosphorylation

Ycf1 phosphorylation was assayed by in-gel phosphoprotein staining. Dephosphorylated Ycf1 for these experiments was generated by treating SEC-purified Ycf1 with Lambda phosphatase (Lambda PP, NEB) for 45 min at 4°C as previously described for CFTR^26^. Following phosphatase treatment, dephosphorylated Ycf1 was subject to a second round of SEC purification to remove excess enzyme immediately prior to use for subsequent biochemical analysis. For in gel phosphoprotein staining, equal quantities (5 ug) of untreated Ycf1 and lambda PP (NEB) treated samples were separated on two separate 10% SDS-PAGE gels, one of which was then stained with Pro-Q Diamond Phosphoprotein Gel Stain (Thermo Scientific) following the manufacture’s protocol to detect protein phosphorylation. The second gel was simultaneously stained with Coomassie Brilliant Blue R-250 as a control for monitoring total protein levels.

### LC-MS/MS analysis of phosphorylated Ycf1

To measure quantitative phosphorylation of purified Ycf1, 10 μg of WT Ycf1 or E1435Q were Trypsin/LysC (Promega, Madison WI) digested with an S-Trap column (ProtiFi, Farmingdale NY) using the manufacturer’s suggested protocol following reduction with DTT and alkylation with IAA with ProteaseMax (0.1% Promega) added to the digestions. The LC-MS/MS analysis was performed using a Q-Exactive Plus (Thermo Fisher Scientific, San Jose, CA) mass spectrometry with an EASY-Spray nanoESI. Peptides were separated using an Acclaim Pepmap 100 trap colum (75 micron ID × 2cm from Themo Scientific) and eluted onto an Acclaim PepMap RSKC analytical column (75 micron ID × 2cm, Thermo Scientific) with a gradient of solvent A (water and 0.1% formic acid) and solvent B (acetonitrile and 0.1% formic acid). The gradients were applied starting with 3-35% Solvent B over 90 minutes, then 25 to 50% solvent B over 20 minutes, 50-95% solvent B over 5 minutes, and a 100% solvent B for 10 min, and then 3% solvent B for 10 min.

Data were collected with a flow rate of 300 nL/min applied with a Dionex Ultimate 3000 RSLCnano system (Thermo Scientific) with data dependent scanning using Xcalibur v 4.0.27.19^62^. A survey scan at 70,000 resolution scanning mass/charge (m/z) 350-1600 with an automatic gain control (AGC) target of 1e^6^ with a maximum injection time (IT) of 65msec, then a high-energy collisional dissociation (HCD) tandem mass spectrometry (MS/MS) at 37 NCE (normalized collision energy) of the 11 highest intensity ions at 17,5000 resolution, 1.5m/z isolation width, 5e^4^ AGC, and 65msec maximum IT. Dynamic exclusion was used to select an m/z exclusion list for 30 sec after single MS/MS and ions with a charge state of +1, 7, >7, unassigned, and isotopes were excluded.

Search of MS and MS/MS data were performed against the Ycf1-E1435Q sequence, the Uniprot *S. cerevisiae* protein database, and a database of common contaminant proteins (including trypsin, keratin – found at ftp://ftp.thegpm.org/fasta/cRAP) with Thermo Proteome Discoverer v 2.4.0.305 (Thermo Fisher Scientific). Fully tryptic peptides with up to 2 missed cleavage sites were considered in MS/MS spectral matches. The variable modifications considered included methionine oxidation (15.995 Da), cysteine carbamidomethylation (57.021 Da), and phosphorylation (79.966 Da) on serine, tyrosine, and threonine. XCorr score cutoffs at 95% confidence were used to identify proteins using a reverse database search^63^. Identification results from proteins and peptides were further analyzed with Scaffold Q+S v 4.11.1 (Proteome Software Inc., Portland OR), which integrates various search results (from Sequest, X!Tandem, MASCOT) and using Bayesian statistics to identify spectra ^64^. We considered protein identification that satisfied the criteria of a minimum of two peptides with 95% confidence levels for protein and peptide.

### Limited proteolysis by trypsin protease

Fixed amounts of Lambda PP treated, and untreated Ycf1 protein (12 μg) sample were incubated for 30 min on ice with trypsin from bovine pancrease (Sigma) at varying concentrations (0, 0.5, 1, 2, 5, 10, and 15 μg/mL). The reaction was stopped by adding 1 μg/mL of soybean trypsin inhibitor (Sigma) and incubating for an additional 15 min on ice. Next, 5 μL of each reaction were mixed with 1X SDS loading dye and 100 mM DTT from each reaction and separated on a 10% SDS-PAGE gel before visualized with Coomassie Brilliant Blue R-250 staining.

## Acknowledgements

We thank the staff at the Life Sciences North Imaging Facility at the University of Arizona and the Pacific Northwest Center for Cryo-EM (PNCC), especially Theo Humphries, Nancy Meyer, and Craig Yoshioko at PNCC for assistance with data collection and helpful advice. We thank the Eyring Materials Center at Arizona State University for assistance with cryo-EM data collection of WT samples. We thank Krishna Parsawar and Cynthia David for their analysis of mass spectrometry-based phosphorylation identification at the Analytical & Biological Mass Spectrometry Facility at the University of Arizona. We also thank Alexei Rohou, Axel Brilot, Tamir Gonen, and Meghan Gupta for helpful discussions with cryo-EM map refinement and model building, and members of the Tomasiak lab and Robert Stroud for helpful discussions and critical reading of this manuscript.

## Author contributions

Conceptualization: TM Tomasiak; Methodology: NKK, TM Thaker, TM Tomasiak; Investigation: NKK, CRM, SZ, SA, DW, TM Thaker, TM Tomasiak; Visualization: NKK, CRM, TM Thaker, TM Tomasiak; Funding acquisition: TM Tomasiak; Project administration: TM Thaker, TM Tomasiak; Supervision: TM Tomasiak; Writing – original draft: NKK, TM Thaker, TM Tomasiak; Writing – review & editing: NKK, CRM, TM Thaker, TM Tomasiak.

## Data and materials availability

All data are available in the main text or the supplementary materials. Structures presented here are available for download from the Protein Data Bank (PDB codes: 7M68 and 7M69) and corresponding EM data from the EMDB (EMD-23690 and EMD-23690, respectively).

