## Supplemental for "The structural basis for regulation of the glutathione transporter Ycf1 by regulatory domain phosphorylation"

**Affiliations:**

### Supplementary Figure 1

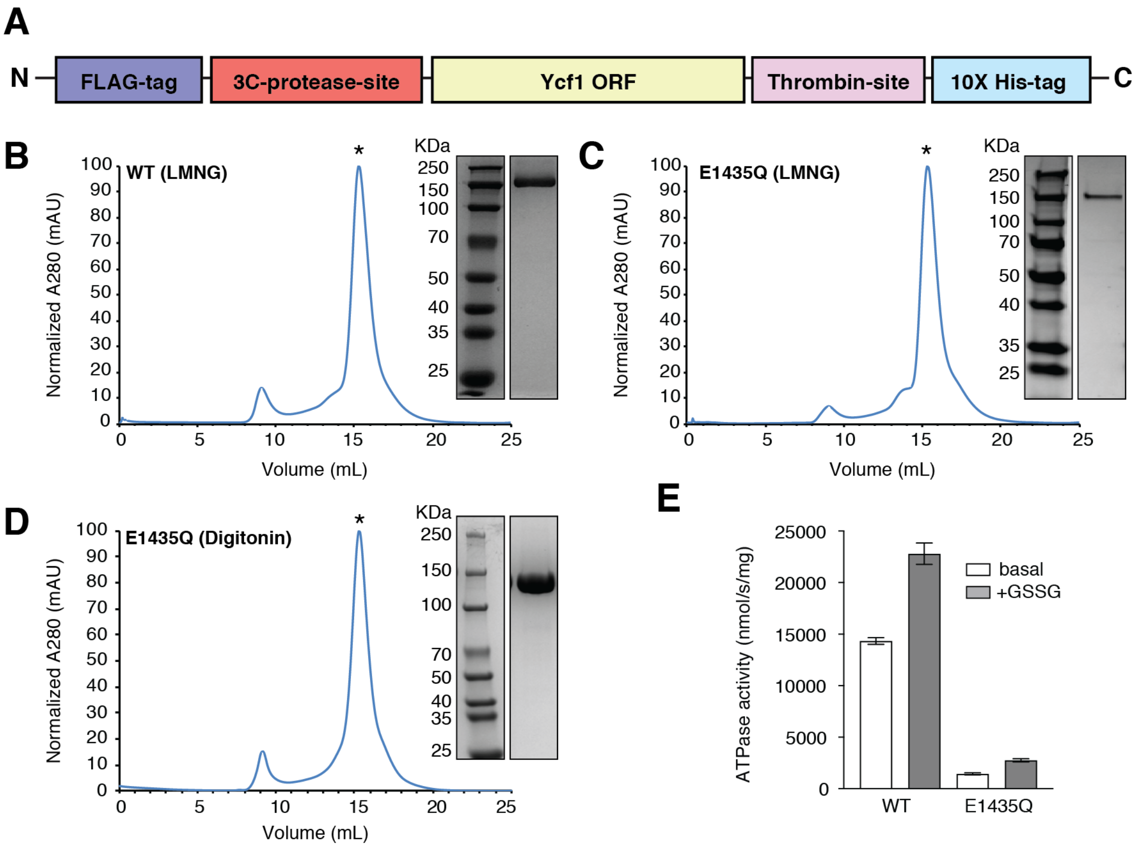

**Fig. S1. Ycf1 purification and biochemical characterization. A.** Construct design for *S. cerevisiae* Ycf1 expression and purification. Representative size exclusion chromatograms (SEC) and corresponding SDS-PAGE results from the purification of (**B**) wild-type (WT) and (**C**) E1435Q Ycf1 in LMNG-containing buffer used for biochemical assays. **D.** SEC profile and corresponding SDS-PAGE result from purifications of E1435Q Ycf1 in digitonin-containing buffer used in the preparation of cryo-EM grids. **E.** Relative ATPase activities in WT and E1435Q Ycf1 in the presence (+GSSG) and absence (basal) of 16 μM oxidized glutathione (GSSG) and 1 mM ATP. Data shown are the mean ± S.D. for n=3 (technical triplicates) and are related to **Main Text Fig. 1F.**

### Supplementary Figure 2

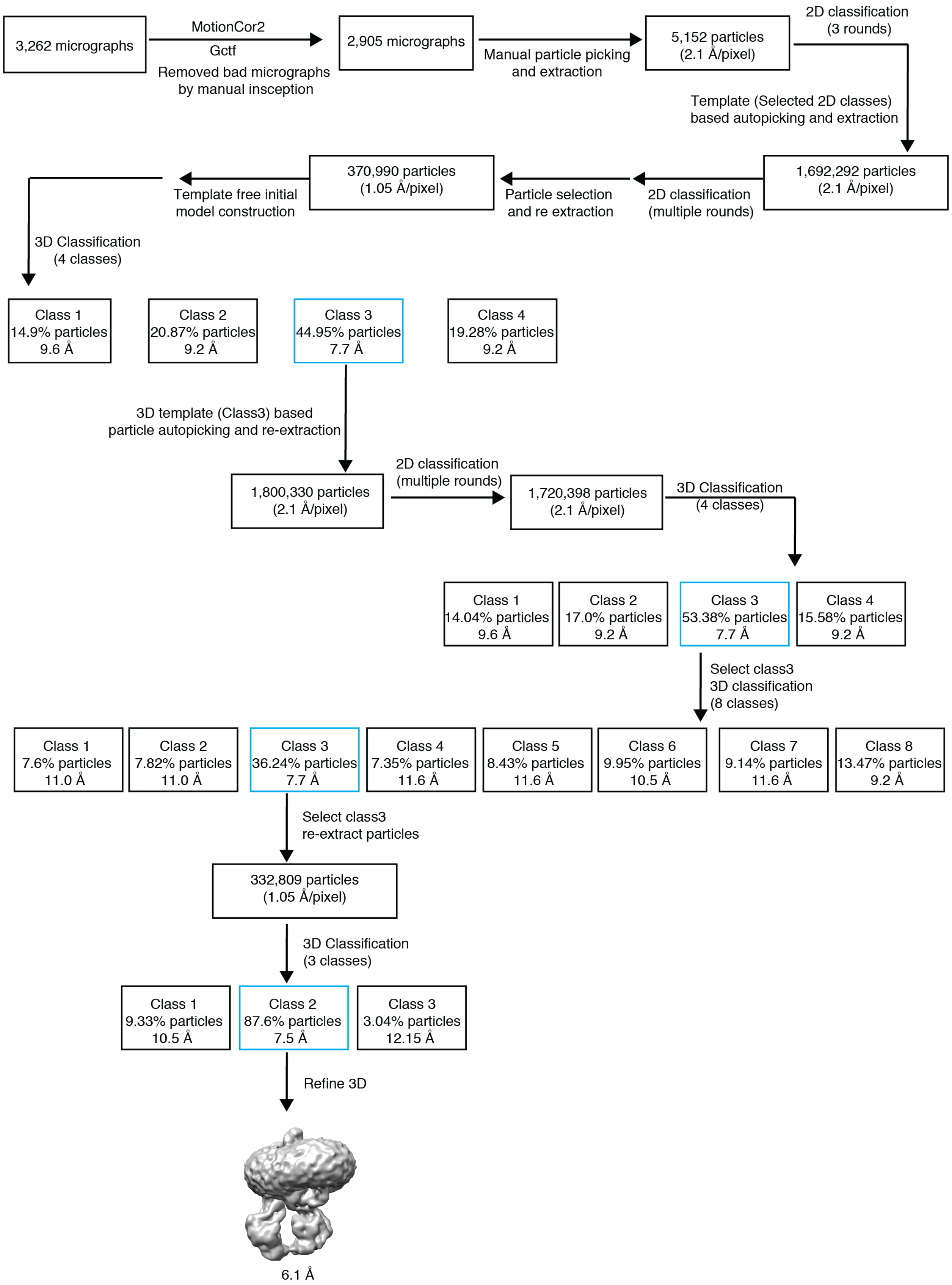

**Fig. S2. Cryo-EM data processing workflow for WT Ycf1.** The image processing pipeline for a dataset of wild-type Ycf1 performed in RELION 3.0. The resulting map was used as a template for particle picking in the E1435Q Ycf1 dataset.

### Supplementary Figure 3

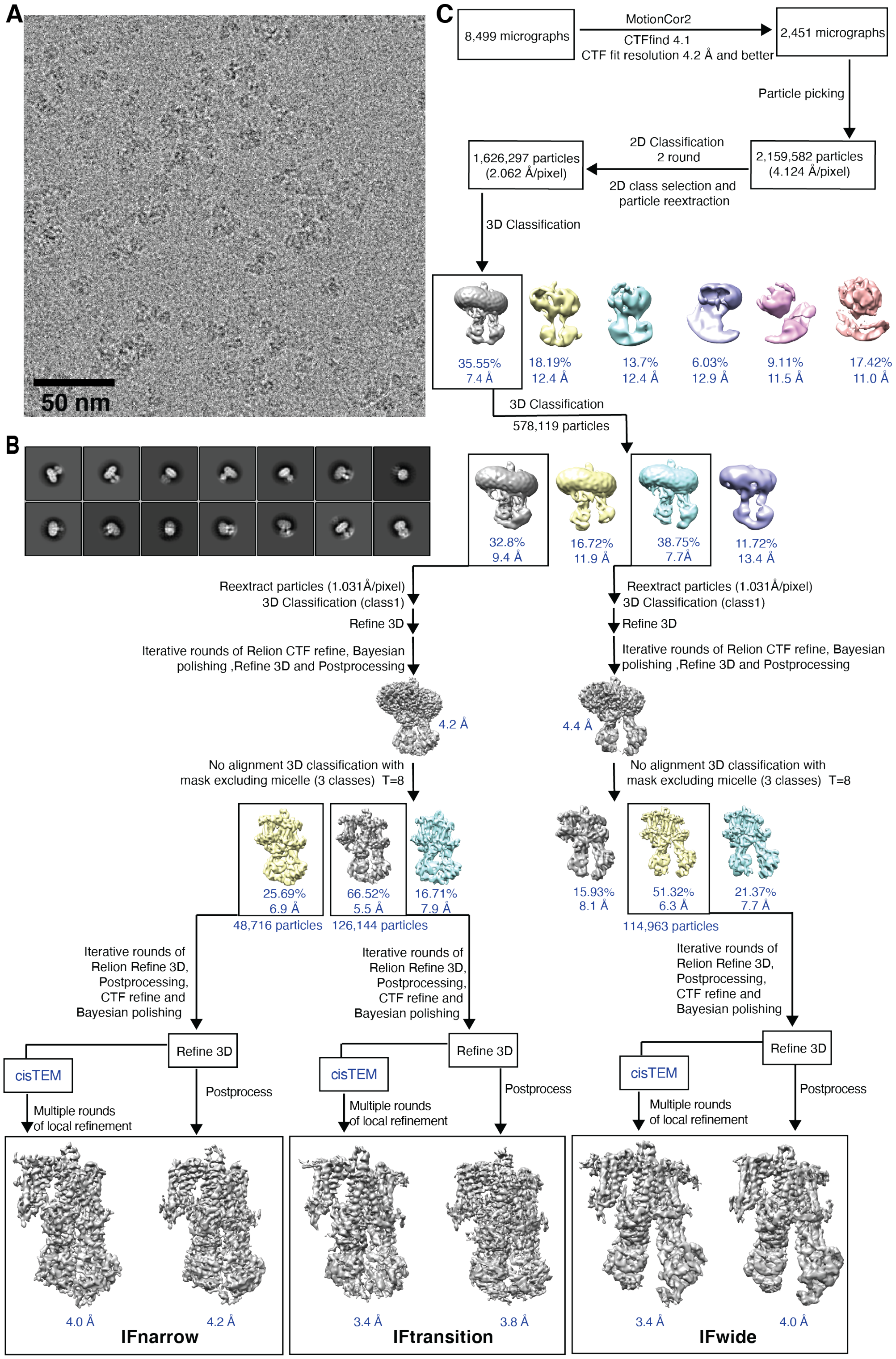

**Fig. S3. Cryo-EM data processing workflow for E1435Q Ycf1. A.** Representative cryo-EM micrographs following motion correction. **B.** Gallery of representative 2D classes for particles used in 3D classification. **C.** Flowchart of map generation and refinement. Initial data were processed in RELION3.1[1]. Initials maps were refined iteratively in RELION3.1 Refine3D. In later rounds refinement was performed using the SIDESPLITTER extension for map reconstruction in RELION3.1 with the “external reconstruct” command[2]. Final rounds of local refinement were performed in cisTEM[3] using particle stacks exported from RELION3.1.

### Supplementary Figure 4

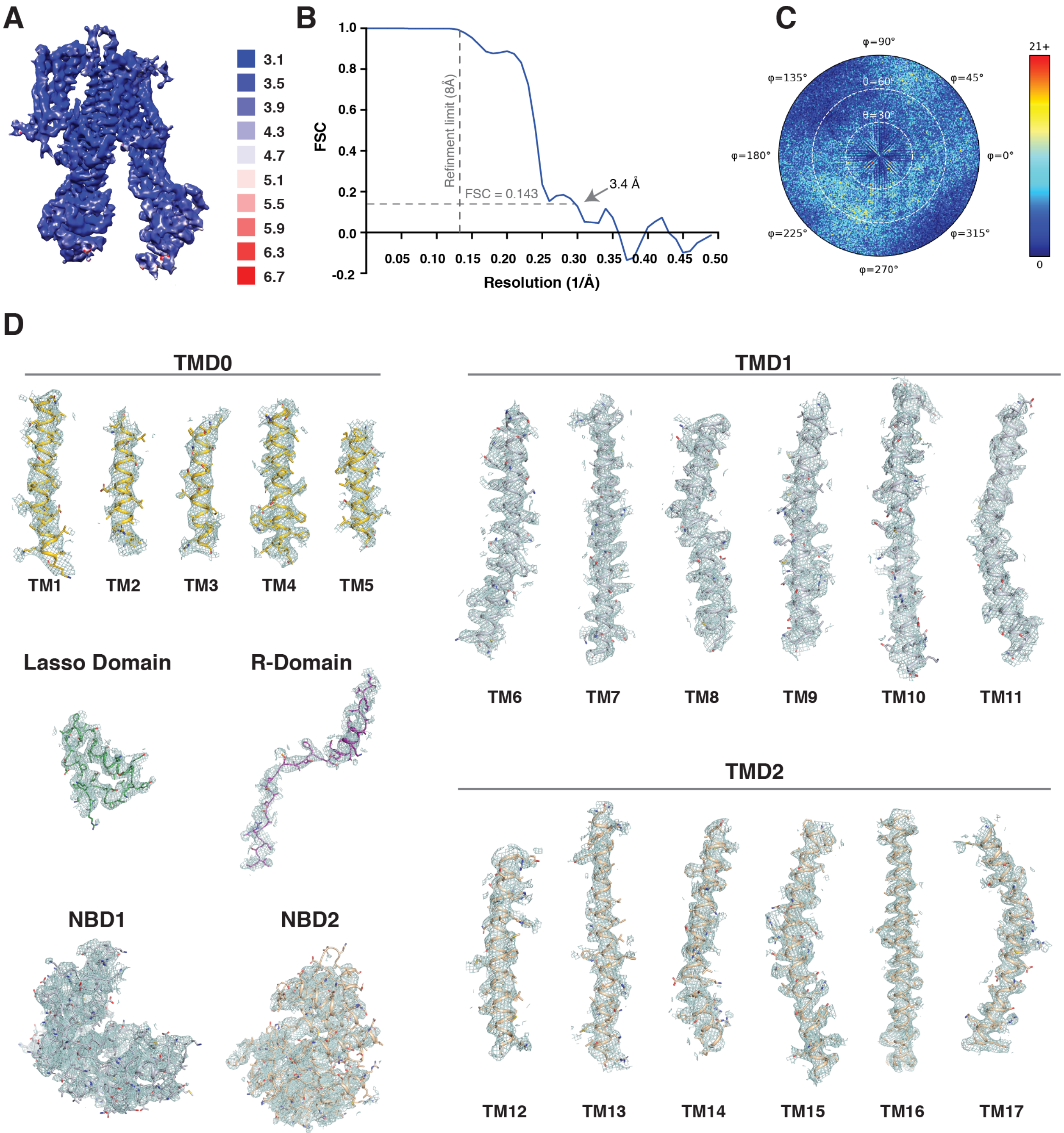

**Fig. S4. Cryo-EM map quality of the Ycf1 IFwide state. A.** Refined map of the E1435Q Ycf1 IFwide conformation colored by local resolution estimated using ResMap[4]. **B.** Fourier Shell Correlation (FSC) plot from cisTEM refinement showing a global resolution of 3.4 Å at a threshold of 0.143. **C.** Angular distribution of particle orientation in the final map reconstruction obtained from CisTEM. **D.** Model and corresponding densities for Ycf1 IFwide domains and transmembrane helices.

### Supplementary Figure 5

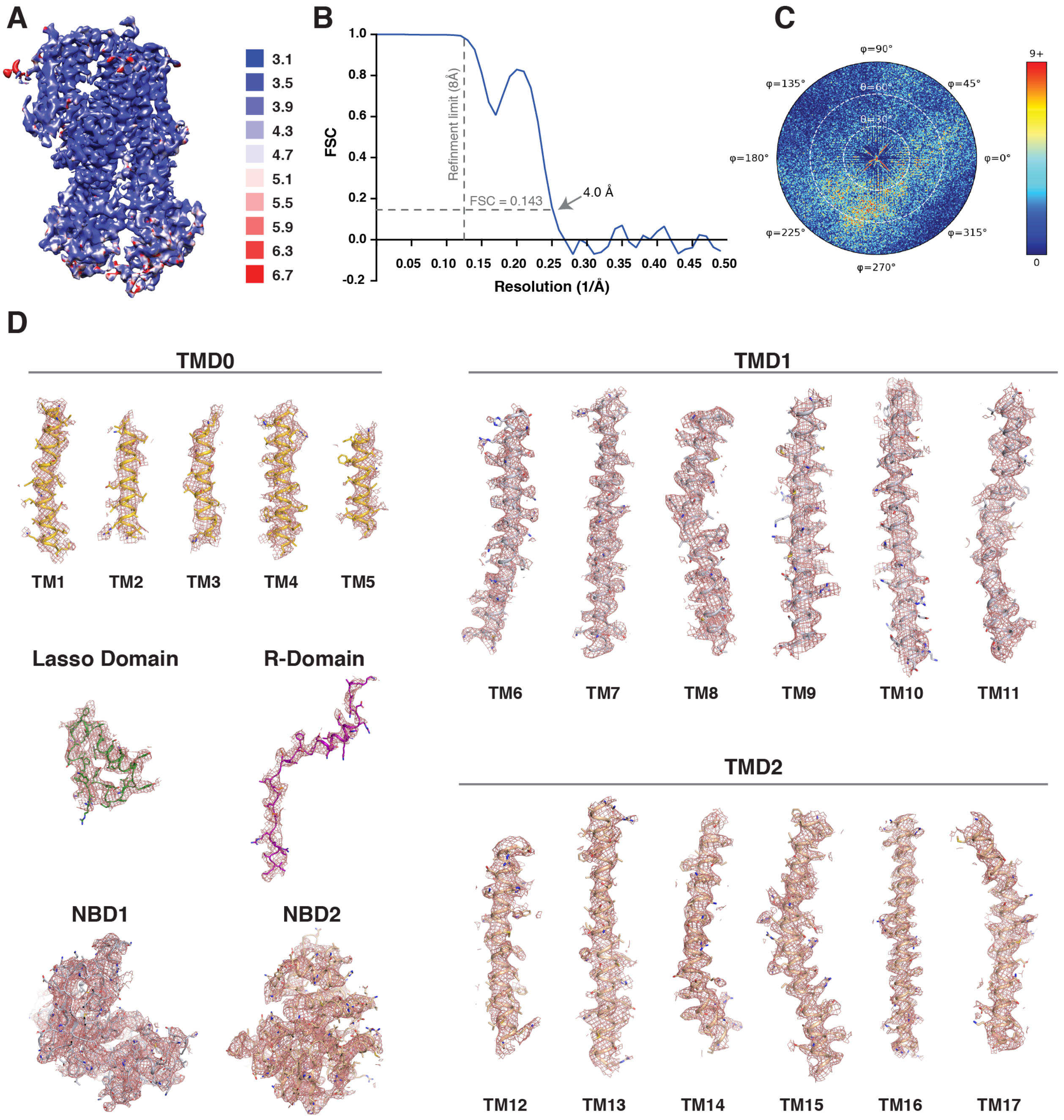

**Fig S5. Cryo-EM map quality of the Ycf1 IFnarrow state. A.** Final refined map of E1435Q Ycf1 IFnarrow conformation. Coloring corresponds to local resolution estimated using ResMap [4]. **B.** Fourier Shell Correlation (FSC) plot from cisTEM refinement showing a global resolution of 4.0 Å at a threshold of 0.143, with corresponding angular distribution of particle orientations in the final map reconstruction obtained from CisTEM shown in (**C**). **D.** Model and corresponding densities for Ycf1 IFnarrow domains and transmembrane helices.

**Supplementary Figure 6**

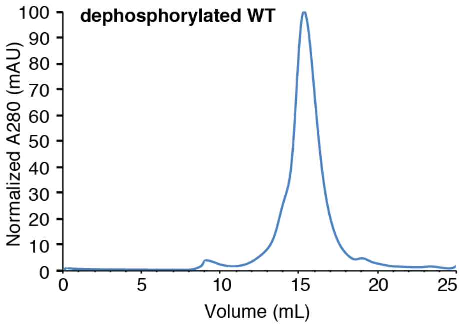

**Fig. S6. Purification of Dephosphorylated Ycf1.** SEC profile for dephosphorylated Ycf1 purified in LMNG-containing buffer following treatment with lambda protein phosphatase.

### Supplementary Figure 7

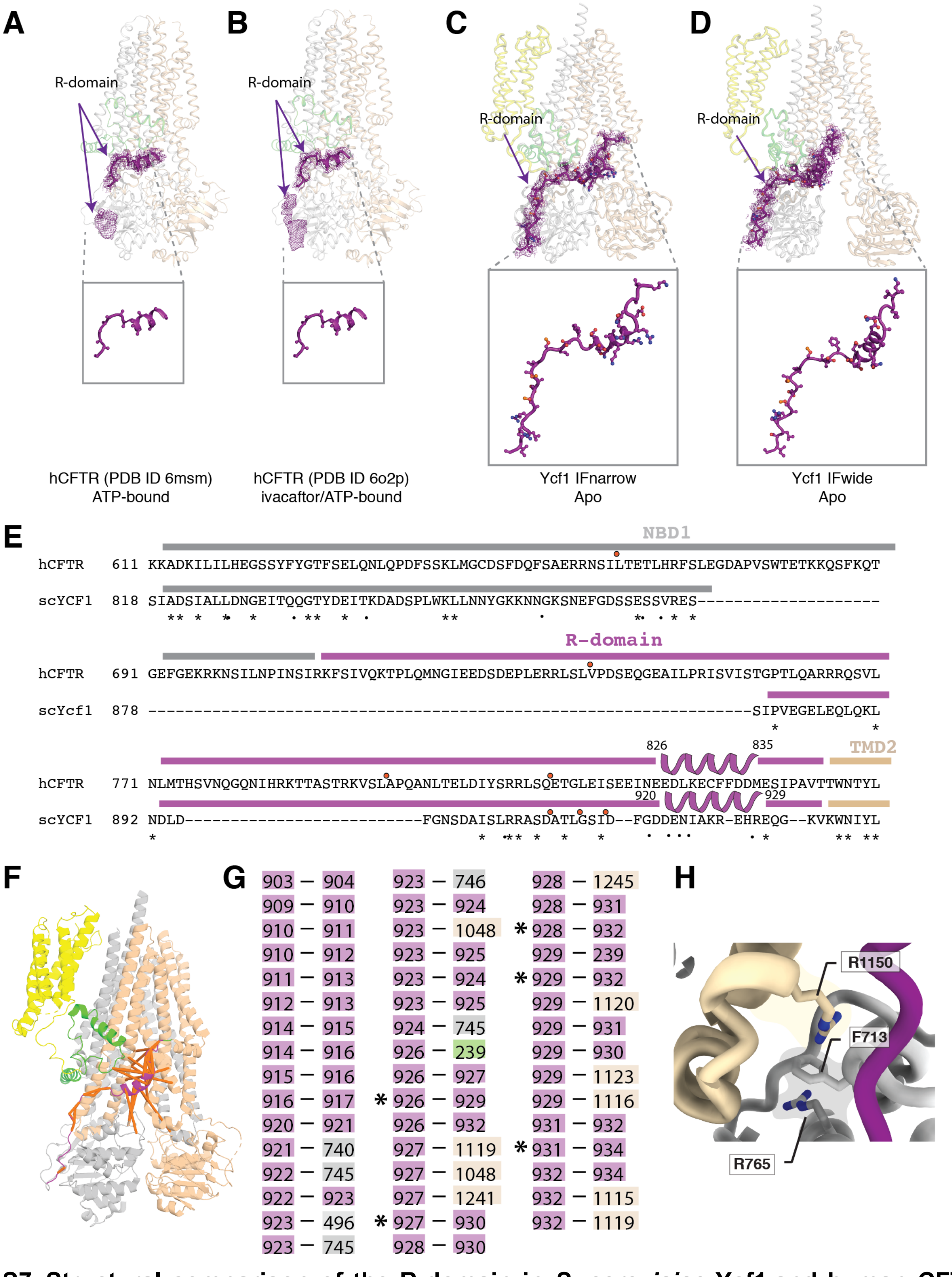

**Fig. S7. Structural comparison of the R-domain in *S. cerevisiae* Ycf1 and human CFTR.** Structures of human CFTR (hCFTR) in the **A.** phosphorylated ATP-bound (PDB ID 6msm[5]) and **B.** ivacaftor- and ATP-bound (PDB ID 6o2p[6]) states showing, in each case, the architecture of the helical portion of the R-domain (cartoon in purple) modeled with polyalanine residues and its corresponding density. The purple mesh represents both assigned and unassigned cryo-EM density, the latter of which the authors also attribute to the R-domain but did not model. **C.** Architecture of the phosphorylated R-domain (purple) in the Ycf1 IFnarrow state and corresponding electron density map into which the model was built (mesh in purple). **D.** The same representation as in (**C**) for the Ycf1 IFwide state. Highlighted below panels (**A-D**) are the amino acid assignments of the R-domain secondary structure in each structure. The lasso domain is shown in green, TMD0 in yellow, TMD1 in light grey, and TMD2 in wheat. **E.** Sequence comparison of the hCFTR and Ycf1 R-domains from an alignment performed using AlignMe[7] and manually adjusted. Residues phosphorylated in both structures are denoted with an orange circle above the corresponding site. Sequence similarity is higher in the C-terminus of the R-domain as compared to the N-terminus. **F.** Network of evolutionary couplings between residues comprising the R-domain (901-935) are shown as orange bands in the structure of Ycf1 IFnarrow colored the same as in (**A-D**). **G.** Annotated list of evolutionary couplings shown in (**F**) between residue pairs colored by the domain in which they reside (same as in **A-D**), with important helical interactions denoted with an asterisk. **H.** Closeup of the interaction network between F713, R1150, R765 in IFnarrow.

**Supplementary Figure 8**

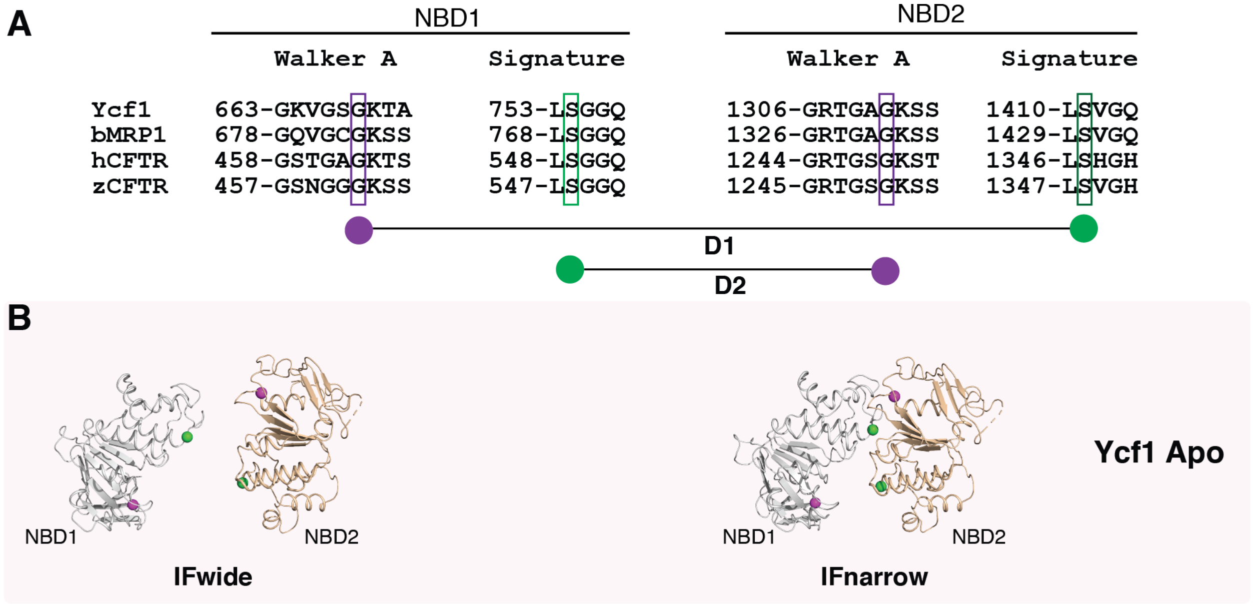

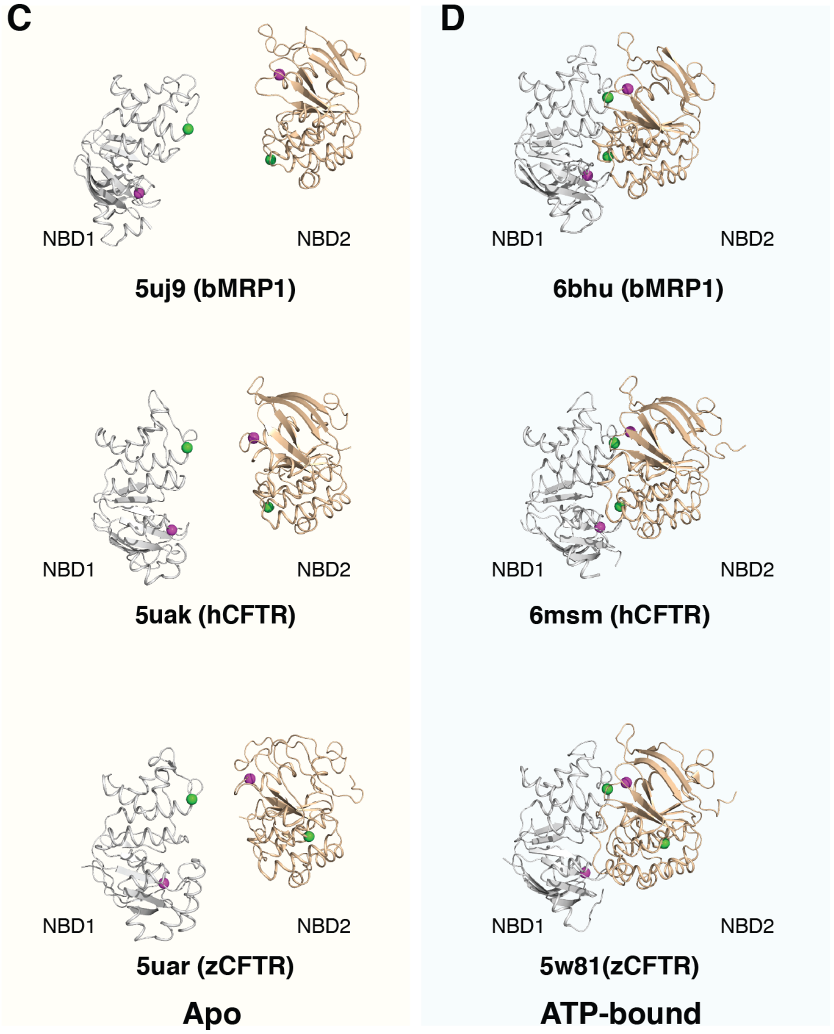

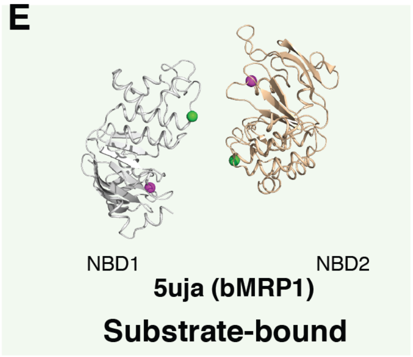

**F**

| Structure | D1 (Å) | D2 (Å) |
|---|---|---|
| Ycf1(IFwide) | 35.4 | 28.8 |
| 5uj9 (bMRP1) | 42.9 | 34.8 |
| 5uak (hCFTR) | 30.6 | 21.0 |
| 5uar (zCFTR) | 47.2 | 19.4 |
| Ycf1(IFnarrow) | 20.9 | 17.8 |
| 5uja (bMRP1) | 27.8 | 24.0 |
| 6bhu (bMRP1) | 8.6 | 9.0 |
| 6msm (hCFTR) | 9.0 | 8.7 |
| 5w81(zCFTR) | 26.0 | 8.6 |

**Fig. S8. Structural comparison of Ycf1 NBD architecture to related C family ABC transporters. A.** Sequence alignment highlighting residues of the Walker A and signature motifs from NBD1 and NBD2 at the NBD dimer interface in Ycf1, bovine MRP1 (bMRP1), human CFTR (hCFTR), and zebrafish CFTR (zCFTR). The interatomic distances between the conserved glycine (purple sphere) of the Walker A motif in NBD1 and conserved serine (green sphere) of the signature motif in NBD2 are denoted as D1. The interatomic distances between the conserved serine (green sphere) of the signature motif in NBD1 and conserved glycine (purple sphere) of the Walker A motif in NBD2 is denoted as D2. **B-E.** Bottom view of NBDs in (**B**) Ycf1 IFwide and IFnarrow structures, (**C**) bMRP1 (PDB ID 5uj9[8], hCFTR (PDB ID 5uak[9]) and zCFTR (PDB ID 5uar[10]) in the apo conformations. (**D**) NBDs bMRP1 (PDB ID 6bhu[11]), hCFTR (PDB ID 6msm[5]) and zCFTR (PDB ID 5w81[12]) in the ATP-bound conformations. **(E**) bMRP1 (PDB ID 5uja[8], in the leukotriene C4 substrate-bound conformation. **F.** Distance measurements between the residues described in (**A**) in the representative ABCC family transporters shown.

**Table S1. Cryo-EM data collection and refinement statistics**

| | | |
|---|---|---|
| **Data collection** | | |
| Microscope | ThermoFisher Titan Krios | |
| Acceleration voltage | 300kV | |
| Detector | Gatan K3 | |
| Image pixel size | 1.031 Å | |
| Defocus range | -0.9 to -2.1 µm | |
| Electron exposure | ~54 electrons /Å$^2$ | |
| Number of frames | 60 | |
| Number of micrographs | 8,499 | |
| **Image Processing** | | |
| | **IFwide** | **IFnarrow** |
| No. of particles in final reconstruction | 114,963 | 48,716 |
| Symmetry | C1 | C1 |
| Final box size (pixels) | 300 | 300 |
| Global resolution (RELION map) | 4.0 Å | 4.2 Å |
| Global resolution (cisTEM map) | 3.4 Å | 4.0 Å |
| FSC threshold | 0.143 | 0.143 |
| Refinement | | |
| Atoms | 21,823 | 21,878 |
| Residues | 1,390 | 1,390 |
| Water | 0 | 0 |
| Supplied Resolution (Å) | 3.4 | 4.0 |
| B-factors (Å$^2$) | | |
| Iso/Aniso (#) | 10887/0 | 10899/0 |
| Protein (min/max/mean) | 87.21/223.07/151.76 | 91.48/221.82/148.98 |
| Bonds (RMSD) | | |
| Length (Å) (# > 4σ) | 0.005 | 0.005 |
| Angles (°) (# > 4σ) | 0.805 | 0.794 |
| **Validation** | | |
| MolProbity score | 1.11 | 1.12 |
| Clash score | 0.92 | 0.78 |
| Ramachandran plot (%) | | |
| Outliers | 0 | 0 |
| Allowed | 4.81 | 5.40 |
| Favored | 95.19 | 94.60 |
| Rotamer outliers (%) | 0.86 | 0.43 |
| **Model vs. Data** | | |
| CC (mask) | 0.70 | 0.70 |
| CC (box) | 0.52 | 0.51 |
| CC (peaks) | 0.34 | 0.32 |
| CC (volume) | 0.70 | 0.71 |
| PDB ID | 7M 69 | 7M68 |

**Table S2.** The quantitative phosphorylation status of wild-type (WT) Ycf1 and Ycf1-E1435Q mutant. Top 5 phosphorylation site with best A score which define the location probabilities.

| | **A-score /Location probability** | |
|---|---|---|
| **Phosphosites** | **WT** | **E1435Q** |
| S908 | 1000 | 1000 |
| T911 | 1000 | 1000 |
| S914 | 1000 | 1000 |
| S903 | 98.4 | 86.37 |
| S251 | 61.94 | 53.98 |
